# The Molecular Signatures Database Revisited: Extending Support for Mouse Data

**DOI:** 10.1101/2022.10.24.513539

**Authors:** Anthony S. Castanza, Jill M. Recla, David Eby, Helga Thorvaldsdóttir, Carol J. Bult, Jill P. Mesirov

**Affiliations:** Department of Medicine, University of California, San Diego, La Jolla, CA 92093, USA; The Jackson Laboratory for Mammalian Genomics, Bar Harbor, ME, 04609, USA; Broad Institute, Cambridge, MA, 02142, USA; Moores Cancer Center, University of California, San Diego, La Jolla, CA 92093, USA

## Abstract

The Molecular Signatures Database (MSigDB) serves as the primary repository of biological signature gene sets for performing Gene Set Enrichment Analysis (GSEA). In the more than 15 years since its creation, MSigDB has served over 290,000 users in their use of GSEA to perform statistically rigorous analysis of coordinated patterns of gene expression changes by leveraging the prior knowledge of tens of thousands of deposited signatures. In that time, the sets provided in MSigDB have been offered exclusively in the human gene space and only minimally supporting analysis of mouse model data through mapping to human genes. Here we present two substantial improvements to MSigDB: first, by providing gene sets from widely used resources in the mouse gene space; and second, by offering substantially improved orthology mapping resources for comparative analysis of both mouse and human datasets.

## INTRODUCTION

The rise of transcriptomic technology, first with microarrays and more recently through sequencing approaches, has fueled a rapid expansion in the availability of molecular-level biological data. These large datasets require interpretation to connect them to meaningful biological and clinical impacts. In 2005, we introduced the Gene Set Enrichment Analysis (GSEA) method with its accompanying Molecular Signatures Database (MSigDB)^1–4^ to enable the identification and estimation of significance of activated biological pathways and processes in molecular data. Briefly, the method takes as input a ranked list of genes derived from an experiment, e.g., by differential expression, and a set of genes that correspond to a biological process or pathway. The method then evaluates if the genes in the gene set are overrepresented at the top or bottom of the ranked list and provides an estimate of significance. When that is the case, the investigator may infer relevance to the underlying biology of the data from which the ranked list is derived, based on the annotations associated with the enriched gene set. Investigators may provide their own gene sets for the analysis, or they can take advantage of the tens of thousands of gene sets available in MSigDB.

The GSEA method is species agnostic, but it does require that the identifiers of the genes in the analyzed data match those of the querying gene sets. However, if there are mismatches, the GSEA application can perform updates and conversions behind the scenes utilizing a gene identifier mapping table. Therefore, a critical piece of supporting infrastructure that MSigDB provides is a set of gene identifier mapping tables in the form of “chip” files that can be used by GSEA to convert the identifiers in a dataset to the gene symbols namespace of a particular version of MSigDB. While originally designed to map microarray probeset ids, chip files are now commonly used to map RNA-seq gene accession IDs, and other gene identifiers such as older versions of gene symbols, etc., to match the MSigDB namespace. These mappings can also be used to provide orthology mapping between model organism genes and human genes, and vice versa.

Historically, the GSEA/MSigDB resource focused on human-specific datasets, offering minimal guidance and support for analysis of model organism data through limited provision of orthology mapping. In recognition of the importance of model organisms, particularly mice, for research into the mechanisms of human disease, we describe here our recent expansion of the MSigDB. This comprises the addition of new procedures to produce substantially improved orthology mapping for mouse datasets and, importantly, the introduction of a new database of mouse-native gene sets. With these developments, we have raised support for mouse data to a “first-class citizen” in the world of enrichment analysis.

## RESULTS

### (1) A New Resource of Mouse-Native Gene Sets

Providing a resource of mouse-native gene sets represents a major expansion of the scope of the MSigDB project. While it is useful for investigators to be able utilize the orthology mapping, we note that some degree of uncertainty exists when discriminating between orthologs and paralogs. In addition, even orthologous genes in human and mouse can have different functions. Elimination of likely functionally divergent genes between mouse and human can be desirable from the perspective of translational research where generalizing biological processes from mouse to human is a goal. However, for research into basic biological processes, the inclusion of all genes, i.e., not excluding species-specific genes, may be key to gaining mechanistic insights and to understanding species-specific biology. Therefore, we invested substantial effort to support analysis in the mouse gene-space through the release of new MSigDB gene sets that were curated from mouse-centric databases and datasets and are specified in native mouse gene identifiers. Thus, they can be used directly with mouse datasets without the need for orthology mapping. The resulting Mouse MSigDB (available at msigdb.org, see Figure 1 and DISCUSSION) leverages the organizational paradigm of the existing human MSigDB and includes the following collections of gene sets designed to operate with natively mouse data:

M1 – A “positional” collection consisting of sets of the genes in each cytogenetic band assembled from the mouse genome version GRCm38 initially and will be updated to GRCm39 as cytogenetic band annotations for this assembly are made available.
M2 – The main collection of molecular pathways consisting of subcollections mined from mouse databases developed by Reactome^5^, WikiPathways^6^, and BioCarta^7^, as well as a subcollection of curated gene sets representing signatures of chemical and genetic perturbations (CGP) derived from mouse-native data from the peer-reviewed literature. The initial release of 932 sets in the M2:CGP collection includes 869 miscellaneous mouse-native gene sets derived from the mouse datasets that were included via orthology conversion in the current human MSigDB C2:CGP collection, signatures of neurodevelopmental identities, and signatures derived from cancer-associated scientific publications reporting results from genetically engineered mice.
M3 – Two subcollections of regulatory target gene sets. One consists of miRNA targets for mouse miRNAs mined from the miRDB project^8^. miRNAs with identical seed sequences were collapsed to one set, and in accordance with miRDB’s recommendations only genes with “high-confidence” target prediction scores (prediction score > 80) were retained for the sets. The other subcollection consists of transcription factor targets derived from mouse ChIP-seq datasets reanalyzed by the Gene Transcription Regulation Database project (GTRD)^9^ to produce high confidence predictions of transcription factor binding sites. We worked with the GTRD team to extract transcription factor binding calls for gene promoter regions, which we loosely defined as the region spanning 1000 base pairs upstream and 100 base pairs downstream of the transcription start site.
M5 – Two subcollections of gene sets derived from ontology hierarchies. One consists of Gene Ontology (GO)^10^ mouse-native gene sets consisting of subcollections of each of the three major GO hierarchies (Biological Process, Cellular Component, and Molecular Function). The other is a collection of 92 gene sets generated from expert curation of the peer-reviewed scientific literature by the Mouse Genome Informatics database (MGI)^11^ team using terms related to cancer biology from the Mammalian Phenotype Ontology (MP)^12^ (Supplementary Table 1). The published sources of the annotations for the genes in these gene sets are available from MGI. The gene sets are refreshed periodically using an automated process as additional genes are annotated with the relevant MP terms.
M8 – Curated cell type markers from single cell data. This collection contains high-quality cell identity signatures from the Descartes Mouse Organogenesis Cell Atlas (MOCA)^13^ which can serve to detect specific cell types in single cell data via enrichment, and gene sets of cell type specific markers of ageing derived from the Tabula Muris Senis or “Mouse Aging Cell Atlas”^14^.
MH – While not derived directly from mouse data, we include an orthology converted version of the MSigDB Hallmarks collection^4^. This enables this popular collection of well-defined biological states and processes to be used on mouse data without requiring orthology mapping back to human and which allows a better comparison of the results using MH with those from appropriate mouse-native collections.

**Figure 1:**
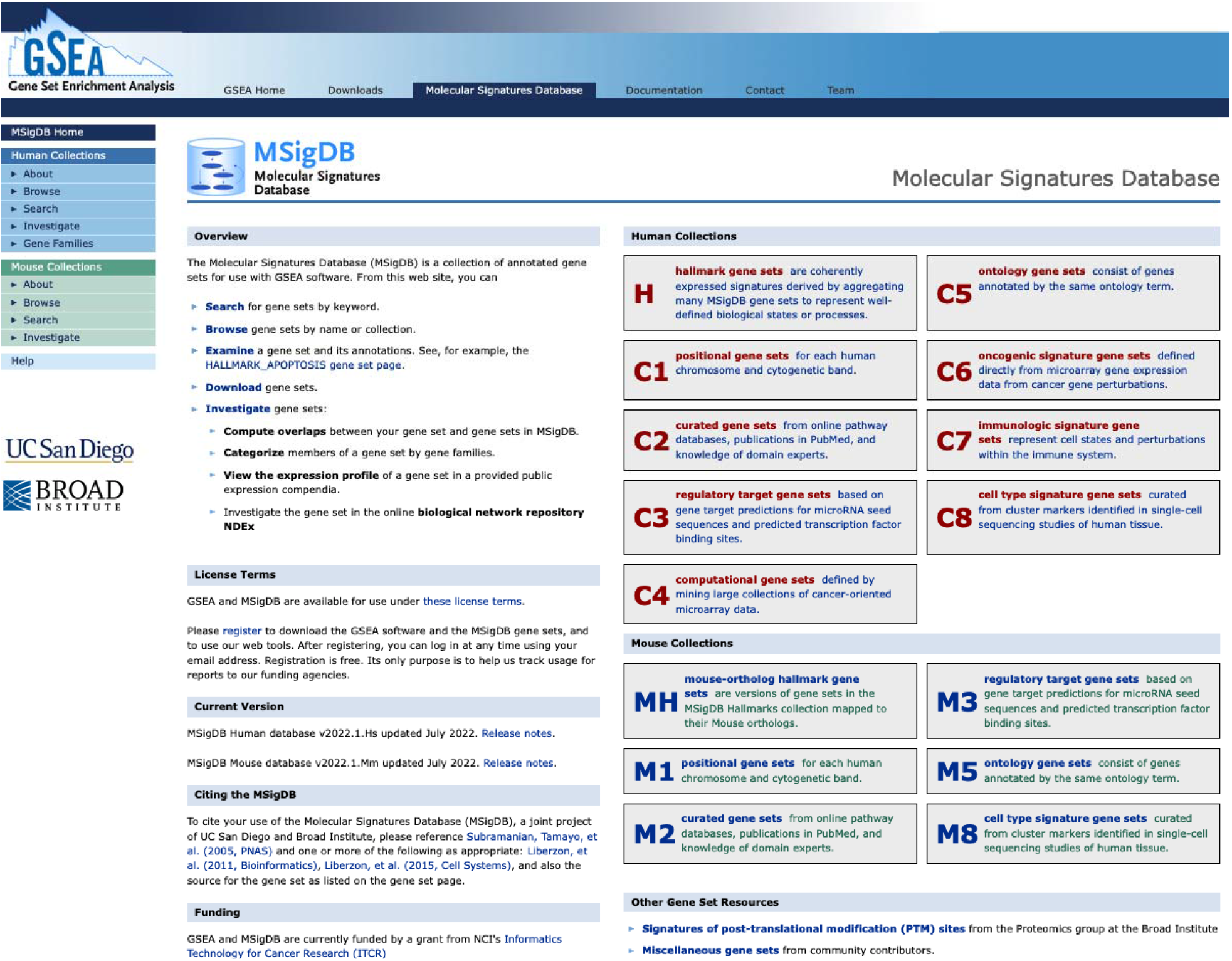
Expanded MSigDB Home Page. The new MSigDB home page (https://msigdb.org) displaying the Human Collections (Top; Red+Blue color scheme) and the Mouse Collections (Bottom; Blue+Green color scheme).

### (2) A New Procedure for Orthology Conversion

For cases where investigators wish to use human gene sets with mouse data or vice versa, we made substantial improvements to MSigDB’s orthology mapping process, taking into account the particular requirements of gene set enrichment analysis. In general, gene orthology annotation approaches catalog all genes that have shared evolutionary relationships. This is done through a myriad of methods, based primarily around comparative sequence analysis, either at the gene or protein level, to estimate the fraction of shared bases in a gene construct, shared structure, regulatory motifs, and gene order conservation. The result of this analysis is a potentially large phylogenetic tree of all known relationships of a gene. This is typically provided as an “all-to-all” mapping table listing all genes with identified orthologous relationships. However, this presents a potential difficulty to cross-species gene set enrichment analysis, as orthologs are likely to have diverged evolutionarily and it may not be clear if the function of the parent gene has been maintained in the paralog(s). This problem is exacerbated when gene duplication events have resulted in multiple paralogs (see Figure 2). In GSEA only a single gene representation of a gene can be present in a dataset. Thus, when multiple orthologs for a given gene cluster are present it is desirable to select the ortholog most likely to preserve the original biological function of the parent gene (the “biologically important relationship” annotated in green in the figure). Unfortunately, it is not always possible to determine an orthologous relationship with high confidence. Indeed, different orthologs may have inherited different amounts of an ancestral gene’s function or undergone complete functional divergence. These multimappings present potential issues for GSEA. A single multi-mapped gene would contribute to the scores of each of these multiple orthologs, some of which may be in the gene set of interest, and some of which may not, thereby potentially simultaneously contributing to, and detracting from, the enrichment score of a set. This can result in unpredictable and undesirable effects on the GSEA enrichment score, as well as potentially increase the score’s contribution from genes that may not be biologically relevant for translational research.

**Figure 2.**
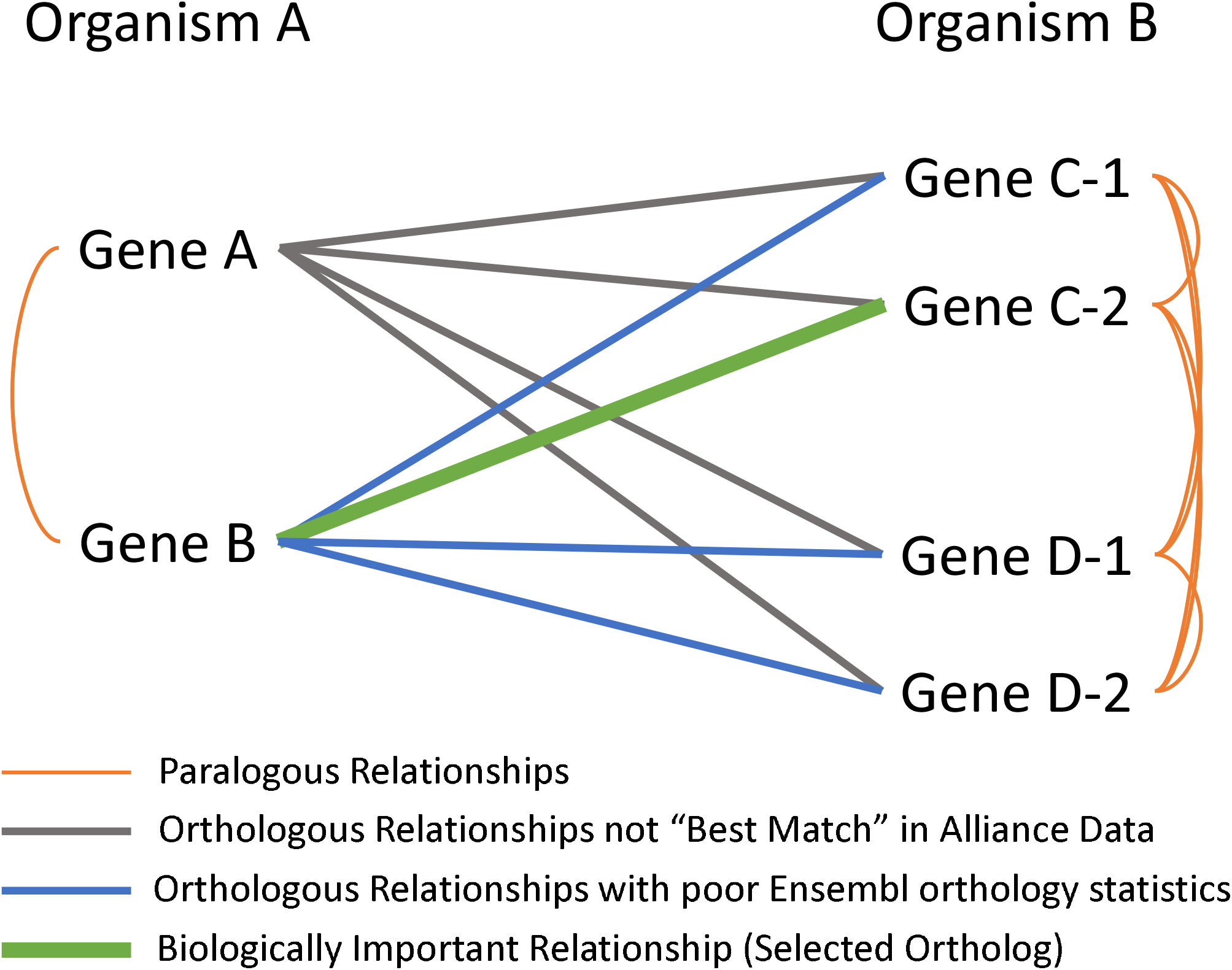
Schematic of Ortholog Relationships. Organism A Gene A and Gene B may be related to Organism B Gene C-1, Gene C-2, Gene D-1, and Gene D-2, but when analyzing data from Organism B how do you determine if a finding from Gene C-2 is relevant for translational work for Organism A, and if so, if it applies to Gene A or Gene B?. Relationships between Gene A and Gene D-1,D-2 are eliminated by not being found as consensus best forward and best reverse orthologs in the Alliance For Genome Resources data. Gene C-2 is selected as the best ortholog for Gene B by scoring higher on whole genome alignment score, gene order conservation score, and identity percentages.

To help mitigate these issues, we adopted a “best match ortholog” approach for producing the mapping tables that underly MSigDB’s “chip” files for orthology identifier conversion between mouse and human, and rat and human. The resulting new procedure is in the form of a two-step process, starting with an initial core set of orthology annotations from the Alliance of Genome Resources^15,16^, which integrates annotations from multiple independent methods via the DRSC (Drosophila RNAi Screening Center) Integrative Ortholog Prediction Tool (DIOPT)^17^, and supplemented with additional orthology data from Ensembl^18,19^. We use the strict orthology assertions for mouse, human, and rat (i.e., annotations that are flagged as “best forward” and “best reverse” ortholog) from the Alliance of Genome Resources data, which excludes many lower confidence matches. Starting from this initial list, we import Ensembl-annotations for these genes for the forward and reverse percentage match, the gene order conservation score, and the whole genome alignment score, for all Ensembl annotated orthologs. These two lists are merged, and Alliance orthologs are filtered further, keeping the gene pairs with the highest whole genome alignment coverage. Any remaining ties are broken using the highest gene order conservation score, highest combined forward, and reverse homology scores, sequentially. The Alliance dataset does not represent a complete accounting of all expressed genes, rather it mainly contains annotated protein coding genes. Additionally, the initial restriction to annotations flagged as “best forward” and “best reverse” orthologs eliminates genes where the methods considered in DIOPT did not come to a consensus on any ortholog match. To prevent introducing a significant gap in the orthology tables if these unmatched genes were excluded, we supplement the Alliance dataset with additional gene information from Ensembl’s orthology annotations. The process described above for importing and filtering Ensembl’s orthology data is then performed for additional ortholog pairs that are present in Ensembl but for which no ortholog was captured in the Alliance dataset.

The resulting improved orthology mapping was used to create new “chip” files for mapping mouse or rat genes to human genes and human or rat genes to mouse genes. These files improve analysis of mouse datasets when used with the human MSigDB, but also allow for the analysis of human datasets with the mouse gene set database. We also used this improved orthology mapping to create the MH collection of Hallmarks described above.

## DISCUSSION

Three main alternative resources currently exist for mouse native gene sets that can be used with GSEA. First, the Gene Set Knowledgebase (GSKB)^20^ offers a number of collections similar in structure to MSigDB. However, the GSKB project has not been updated since 2012, resulting in the main collections of interest being significantly out of date. Second, the EnrichR project^21^ offers a number of “Mouse” gene set “libraries”. However, the files provided for these libraries consist of gene symbols in the human standardized naming convention, not mouse, and do not provide any details about the namespace, such as symbol versions or sources. Third, the Bader Lab presents, and frequently updates, gene sets (http://baderlab.org/GeneSets/) in the mouse namespace. However, rather than representing a true repository of mouse-native gene sets, the sets provided in this resource are almost all annotated as “translated from Human using Homologene”, i.e., they are not truly mouse native gene sets. Furthermore, none of these three resources provide any identifier mapping tables, which are necessary for the conversion required for a GSEA analysis when the gene identifiers of the dataset are not the same as the gene sets. Finally, GO2MSig^22^ is an additional resource that previously provided scripts and gene sets derived from the GO database for many species including mouse, but this data is no longer available. We note however, that we consulted on the development of the GO2MSig scripts, and we have since adapted them to improve the process of producing the MSigDB GO collections.

Although the Mouse MSigDB is a new resource, it benefits from the maturity of the existing human MSigDB and accompanying website. When preparing the new mouse gene sets, we followed the same procedure that we have honed over the years for our human gene sets. Sets are collected from their respective sources with the lowest-level identifiers available, i.e., Ensembl Gene IDs, NCBI Gene IDs, MGI Gene IDs, etc. These identifiers are preferred over gene symbols for the process of curation as the ID’s association to the underlying gene locus or model is typically stable over time, whereas symbols are commonly revised to reflect changes in functional understanding^23^. These low-level identifiers are then mapped to a consistent and reproducible version of gene symbols. In particular, we use data retrieved from Ensembl’s BioMaRt service^24,25^ to map the identifiers to the latest version of the Ensembl transcriptome assembly, and the symbols captured by Ensembl in that snapshot, available at the time of each MSigDB release. Sets derived from some hierarchical resources, such as Reactome and Gene Ontology, also undergo a redundancy filtering procedure to select a single representative set from groups of identical or highly similar candidate sets. This helps prevent FDR inflation in GSEA results, which can be caused by multiple identically, or nearly identically scoring gene sets. Briefly, when creating the sets from each of these resources, we compute Jaccard coefficients for all pairs of candidate sets, selecting a Jaccard coefficient greater than 0.85 as “highly similar”. We then cluster highly similar sets into groups and apply two rounds of filtering for each group. First, we keep the largest set in the group and discard the smaller sets. Then, in the case of identically sized sets, the set closest to the hierarchy root (i.e., the most general term) is retained. This method was adapted from our prior work generating the MSigDB Hallmarks collection^4^.

As an assessment of the performance of the orthology mapping procedure compared to providing gene sets in the mouse-native space, we analyzed the 869 mouse-derived gene sets that had been provided in an orthology mapped form in the human C2:CGP collection, and the corresponding gene sets in their native gene-space in the mouse M2:CGP collection (see the description of the M2 collection in RESULTS). For both databases, the gene set creation involved mapping the mouse identifiers in the original source data for these sets (e.g., RNA-seq accessions, microarray probe IDs, or gene symbols from across decades of research) to consistent gene symbols, either human (via orthology mapping) or mouse. When mapping these identifiers to gene symbols, regardless of the target species, some number of identifiers generally do not map. This can be for many reasons – mapping to a construct that is not recognized by both Ensembl and NCBI genome assemblies, mapping to a gene symbol that has been withdrawn, etc. From this data we derive a “mapping success rate” or “mapping rate”, i.e., the number of gene IDs successfully converted to gene symbols. This mapping rate varied substantially from gene set to gene set, with 75% of sets losing less than 5% of their members through the orthology conversion (Figure 3). Of the 869 sets, 301 sets resulted in an identical number of members in the human and mouse databases. In these sets, on average 3.6 of a set’s original identifiers did not map to an approved gene symbol, neither mouse nor human. 563 sets had higher mapping rates in their native mouse gene space for the mouse database. These improved sets contained approximately 5.4% more final mapped genes in the mouse database than in the orthology converted version in the current human database. This translates to an average of 6.5 genes that were gained when these sets were provided in the native mouse database, although much larger numbers of affected genes were observed in a small number of these sets. For example, in one of the larger sets (1128 original identifiers), 98 genes were gained, from 900 mapped genes in the orthology converted form in the human database to 998 genes in the mouse database (8.7%), and 50 genes (21%) in another set (236 original identifiers mapped to 173 genes orthology converted to human and 223 genes mapped to mouse). In the total 869 sets, an overall mapping rate improvement of 3.6% was found in the mouse database. Interestingly, and contrary to our expectations, 5 sets were observed where the mapping rate was better in the orthology conversion. The reason for this improvement is unclear, however one possibility is the restriction of the final MSigDB gene space to include only genes that have a recognized construct in both Ensembl and NCBI, while the orthology conversion process does not require an assigned NCBI gene ID. In conclusion, the native mouse gene sets in the M2 collection generally resulted in moderately improved gene set fidelity compared to the orthology-converted sets in the human database. However, the performance of the orthology tables – 96.4% mapping rate on average – indicates that while not strictly as good as native sets, orthology-converted data is still likely to provide reasonable results in most cases.

**Figure 3:**
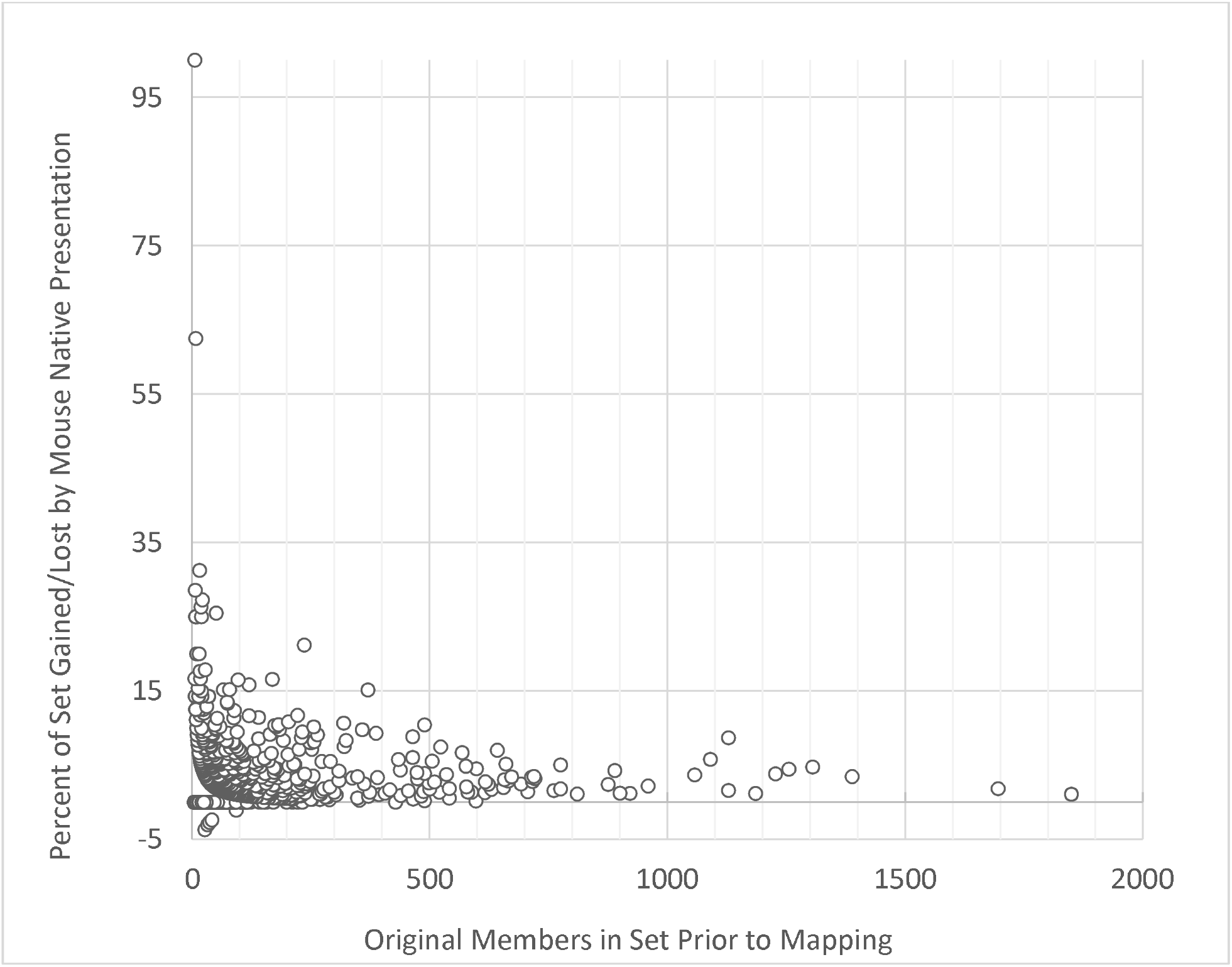
Comparison of mapping rates for mouse gene sets previously orthology converted to human vs the same gene sets in MSigDB’s native mouse database. Positive values along the y-axis indicate that a gene set exhibited a higher mapping rate to valid gene symbols in the native (mouse) species than when the set was mapped to human orthologs. Likewise, a negative y-axis value indicates that the set exhibited decreased mapping rate in the native (mouse) species (i.e. a greater mapping rate upon orthology conversion to human.)

In summary, here we have described two major improvements to the Molecular Signatures Database that offer substantial increased functionality to the popular GSEA method and other enrichment methods and tools that utilize the MSigDB resource. First, we made available a new series of gene set collections that enable the native use of mouse-derived datasets in GSEA analyses. The initial release of Mouse MSigDB delivers a substantial offering of gene sets, including collections of sets that parallel some of the most popular collections in the human MSigDB. We continue to develop Mouse MSigDB and the collections will be expanded further in future releases. Second, we described a redesigned orthology conversion process that improves consistency and reliability. By offering support for native analysis in mouse and for mapping reciprocally from mouse data to use with human sets and from human data to use with the new mouse-native sets, these enhancements represent a significant step forward in the potential translational impact of GSEA results and put the mouse and human MSigDB collections on equal footing.

## References

1. Mootha, V. K. et al. PGC-1α-responsive genes involved in oxidative phosphorylation are coordinately downregulated in human diabetes. Nat. Genet. (2003). doi:10.1038/ng1180

2. Subramanian, A. et al. Gene set enrichment analysis: A knowledge-based approach for interpreting genome-wide expression profiles. Proc. Natl. Acad. Sci. U. S. A. (2005). doi:10.1073/pnas.0506580102

3. Liberzon, A. et al. Molecular signatures database (MSigDB) 3.0. Bioinformatics (2011). doi:10.1093/bioinformatics/btr260

4. Liberzon, A. et al. The Molecular Signatures Database Hallmark Gene Set Collection. Cell Syst. (2015). doi:10.1016/j.cels.2015.12.004

5. Gillespie, M. et al. The reactome pathway knowledgebase 2022. Nucleic Acids Res. (2022). doi:10.1093/nar/gkab1028

6. Martens, M. et al. WikiPathways: Connecting communities. Nucleic Acids Res. (2021). doi:10.1093/nar/gkaa1024

7. Nishimura, D. BioCarta. Biotech Softw. Internet Rep. (2001). doi:10.1089/152791601750294344

8. Chen, Y. & Wang, X. MiRDB: An online database for prediction of functional microRNA targets. Nucleic Acids Res. (2020). doi:10.1093/nar/gkz757

9. Kolmykov, S. et al. Gtrd: An integrated view of transcription regulation. Nucleic Acids Research (2021). doi:10.1093/nar/gkaa1057

10. Carbon, S. et al. AmiGO: Online access to ontology and annotation data. Bioinformatics (2009). doi:10.1093/bioinformatics/btn615

11. Ringwald, M. et al. Mouse Genome Informatics (MGI): latest news from MGD and GXD. Mamm. Genome (2022). doi:10.1007/s00335-021-09921-0

12. Smith, C. L. & Eppig, J. T. The mammalian phenotype ontology: Enabling robust annotation and comparative analysis. Wiley Interdiscip. Rev. Syst. Biol. Med. (2009). doi:10.1002/wsbm.44

13. Cao, J. et al. The single-cell transcriptional landscape of mammalian organogenesis. Nature (2019). doi:10.1038/s41586-019-0969-x

14. Almanzar, N. et al. A single-cell transcriptomic atlas characterizes ageing tissues in the mouse. Nature (2020). doi:10.1038/s41586-020-2496-1

15. Bult, C. J. et al. The alliance of genome resources: Building a modern data ecosystem for model organism databases. Genetics (2019). doi:10.1534/genetics.119.302523

16. Agapite, J. et al. Alliance of Genome Resources Portal: Unified model organism research platform. Nucleic Acids Res. (2020). doi:10.1093/nar/gkz813

17. Hu, Y. et al. An integrative approach to ortholog prediction for disease-focused and other functional studies. BMC Bioinformatics (2011). doi:10.1186/1471-2105-12-357

18. Herrero, J. et al. Ensembl comparative genomics resources. Database (2016). doi:10.1093/database/bav096

19. Pignatelli, M. et al. NcRNA orthologies in the vertebrate lineage. Database (2016). doi:10.1093/database/bav127

20. Lai, L. et al. GSKB: A gene set database for pathway analysis in mouse. bioRxiv (2016).

21. Kuleshov, M. V. et al. Enrichr: a comprehensive gene set enrichment analysis web server 2016 update. Nucleic Acids Res. (2016). doi:10.1093/nar/gkw377

22. Powell, J. A. C. GO2MSIG, an automated GO based multi-species gene set generator for gene set enrichment analysis. BMC Bioinformatics (2014). doi:10.1186/1471-2105-15-146

23. Bruford, E. A. et al. Guidelines for human gene nomenclature. Nature Genetics (2020). doi:10.1038/s41588-020-0669-3

24. Durinck, S., Spellman, P. T., Birney, E. & Huber, W. Mapping identifiers for the integration of genomic datasets with the R/ Bioconductor package biomaRt. Nat. Protoc. (2009). doi:10.1038/nprot.2009.97

25. Kinsella, R. J. et al. Ensembl BioMarts: A hub for data retrieval across taxonomic space. Database (2011). doi:10.1093/database/bar030

